# Selective attention to sound features mediates cross-modal activation of visual cortices

**DOI:** 10.1101/855882

**Authors:** Chrysa Retsa, Pawel J. Matusz, Jan W.H. Schnupp, Micah M. Murray

## Abstract

Contemporary schemas of brain organization now include multisensory processes both in low-level cortices as well as at early stages of stimulus processing. Evidence has also accumulated showing that unisensory stimulus processing can result in cross-modal effects. For example, task-irrelevant and lateralized sounds can activate visual cortices; a phenomenon referred to as the auditory-evoked contralateral occipital positivity (ACOP). Some claim this is an example of automatic attentional capture in visual cortices. Other results, however, indicate that context may play a determinant role. Here, we investigated whether selective attention to spatial features of sounds is a determining factor in eliciting the ACOP. We recorded high-density auditory evoked potentials (AEPs) while participants selectively attended and discriminated sounds according to four possible stimulus attributes: location, pitch, speaker identity or syllable. Sound acoustics were held constant, and their location was always equiprobable (50% left, 50% right). The only manipulation was to which sound dimension participants attended. We analysed the AEP data from healthy participants within an electrical neuroimaging framework. The presence of sound-elicited activations of visual cortices depended on the to-be-discriminated, goal-based dimension. The ACOP was elicited only when participants were required to discriminate sound location, but not when they attended to any of the non-spatial features. These results provide a further indication that the ACOP is not automatic. Moreover, our findings showcase the interplay between task-relevance and spatial (un)predictability in determining the presence of the cross-modal activation of visual cortices.

## Introduction

Cross-modal modulations of activity within visual cortices by auditory stimuli have been reported in multiple brain regions of the visual system and at diverse latencies after sound presentation (reviewed in Murray et al., 2016 and De Meo et al., 2015). Sounds can modulate responses in visual cortex and affect early sensory and perceptual processing (e.g. Murray et al., 2016; Hillyard et al., 2016;Mercier et al., 2013; Cate et al., 2009). Multisensory integration of information as well as orienting of spatial attention towards the location of the sound both contribute to these effects. It has been shown that sounds: i) improve the detection and discrimination of concurrently presented visual stimuli (e.g. McDonald et al., 2000; Gleiss and Kayser, 2014; Kayser and Kayser, 2018), ii) increase their subjective intensity (Stormer et al., 2009; Odgaard et al., 2004; Lovelace et al., 2003), iii) speed up responses to visual targets and enhance their attentional selection (Brang et al., 2015; Cappe et al., 2010; Matusz & Eimer, 2011, 2013; Matusz et al. 2015a; 2019a, 2019b; also Kayser et al., 2017), and iv) even contribute to visual perceptual filling-in (Tivadar et al., 2018).

Such cross-modal activations in the visual cortex by auditory stimuli have been found to start as early as ~40-50ms (e.g. Giard and Peronnet, 1999; Raij et al., 2000; Raij et al., 2010; Molholm et al., 2002; Cappe et al., 2010). In addition, in a series of TMS studies, Romei et al. 2007, 2009, 2013; Spierer 2013, showed that auditory stimuli enhance the excitability of primary visual cortex as assayed via phosphene induction. More recently, it was demonstrated by an electrocorticography study that laterally presented sounds evoked primary visual cortex responses at both early (28-100ms) and later (200-500ms) time windows (Brang et al., 2015; Mercier et al., 2013). Interestingly, this cross-modal activation was driven by contralateral sounds and was observed maximally at anterior calcarine sites, suggesting it is spatially specific (Leo et al., 2011; Bolognini et al. 2013). These visual cross-modal activations are potentially mediated by direct anatomical connections between primary auditory and primary visual cortices in animals (Falchier et al., 2010; 2002; Rockland & Ojima, 2003; Clavagnier et al., 2004; Budinger et al., 2006) and in humans (Beer et al., 2011; Eckert et al., 2008).

The conditions under which visual cross-modal activations occur as well as their functional significance remains unclear. There is some evidence indicating that they are task-dependent and category specific (e.g. Campus et al., 2017; reviewed in Ten Oever et al., 2016 and De Meo et al., 2015). Recent event-related potential (ERP) studies have shown that laterally-presented sounds activate contralateral extrastriate visual cortices (McDonald et al., 2013; Feng et al., 2014; Hillyard et al., 2016). The auditory-evoked contralateral occipital positivity (ACOP) is the ERP marker of this cross-modal activation, which starts at approximately 250ms post-stimulus and is localised in the ventrolateral extrastriate visual cortex. The ACOP was initially observed to be elicited by lateralised auditory cues that preceded either visual or auditory targets. In these original experiments, the sound was spatially and temporally unpredictable. Moreover, the magnitude of the ACOP has been linked to better visual perceptual performance (McDonald et al., 2013; Feng et al., 2014). Notably, the ACOP can be elicited even in a context involving passive listening, and in the absence of task-relevant visual stimuli (Matusz et al., 2016). However, in this passive-task paradigm, the ACOP critically depended on the location of the lateralised sounds being unpredictable. These results suggest that the ACOP is a context-contingent process (Matusz et al., 2016).

In all previous experiments reporting the ACOP, the sounds that elicited it were task-irrelevant. It is unknown if cross-modal activation of visual cortices by sounds, as indexed by the ACOP, is impacted by selective attention to specific task-relevant acoustic features. Addressing this question was the primary focus of the present study where we queried whether all attended, but spatially unpredictable, sounds can induce the ACOP. In all of the previous studies on the ACOP mentioned above, the task context involved spatial location either as a task-relevant dimension or, in the case of passive listening, implicitly involved a spatial dimension in the paradigm with to-be-ignored distractors. As such, it is possible that the ACOP is elicited only when spatial location is relevant to the currently performed task. In order to verify this hypothesis, we instructed participants to complete a series of two-alternative-forced-choice tasks involving sounds varying across four perceptual dimensions: location (left/right), pitch (high/low), syllable type (“ti”/”ta”) or speaker identity (man/boy) (Retsa et al., 2018). Across blocks of trials, we explicitly asked participants’ to attend to and discriminate one of these specific features of the sounds. All four discrimination tasks involved identical sounds. The sounds themselves and required tasks were closer to real-world situations compared to the simple beeps used in previous work, thus arguably making the study more ecologically relevant (see Matusz et al. 2019c for more in-depth discussion of how to increase ecological validity of theories of selective attention).

Therefore, we investigated whether task-relevant sounds can trigger the ACOP and whether its strength and even presence or absence depends critically on the task-relevant sound feature. By using an active auditory task, we further explored to what extent and under which circumstances visual cortices are recruited during and perhaps contribute to auditory discrimination. We employed three different object-related tasks, in order to verify whether the ACOP is sensitive to some but not other type of sound features, such as simple tones, as opposed to more naturalistic sounds, such as syllables. Previous studies of ACOP have identified the *sufficient* conditions for the ACOP to manifest (task-irrelevance and spatial unpredictability of the sounds). In the present study, we sought to determine some of the *necessary* conditions for cross-modal activation of visual cortices by laterally presented sounds. We hypothesised that the ACOP is critically dependent on the spatial location being relevant to the currently performed task.

## Materials and methods

### Participants

Sixteen healthy unpaid volunteers (nine female; aged 24–49 years; mean±SD = 28±5 years) provided informed consent to participate in the experiment. All procedures were approved by the Cantonal Ethics Committee. Fifteen of the participants were right-handed and one was left-handed, as assessed with the Edinburgh questionnaire (Oldfield, 1971). As the ERP analyses focused on contralateral versus ipsilateral differences in brain responses (after collapsing ERPs elicited by left-sided and right-sided sounds), these differences should not be influenced by participants’ handedness. None of the subjects reported current or prior neurological or psychiatric illnesses. All participants reported normal hearing and had normal or corrected-to-normal vision. Aspects of this dataset were previously published in a study focusing on parallel processing of object-related feature dimensions in the auditory system (Retsa et al., 2018). However, in that work no analyses based on the lateralisation of the stimuli were performed.

### Apparatus and Stimuli

The participants were seated at the centre of a sound-attenuated chamber (whisper room model 102126 E) and acoustic stimuli were delivered over insert earphones (Etymotic model ER-4P; www.etymotic.com) at a sampling rate of 48kHz. Stimulus intensity was approximately 75dB SPL at the ear. All sounds had duration of 550ms.

Auditory stimuli varied in four dimensions (two levels per dimension: syllable, relative pitch, speaker identity, perceived location), resulting in 16 stimuli. The auditory stimuli were generated by systematically changing a sample of vocalizations from a syllable set recorded by the Cambridge Centre for the Neural Basis of Hearing (CNBC). The original recordings were kindly provided by Prof. Roy Patterson (see Ives, et al., 2005 for details). We chose the syllables /ta/ and /ti/, from the original syllable set, which had been spoken by a single male adult speaker in a quiet room recorded with a Shure SM58-LCE microphone and digitized at 48 kHz. From these recordings, we generated natural-sounding morphs of the /ta/ and /ti/ syllables that were identical in sound intensity and duration and differed only in systematic shifts of their harmonic and formant frequencies to create high-pitched or low-pitched versions in the voice of a man or a boy respectively. To generate the “baritone” voice typical of a large adult man, the formant frequencies of the syllable were scaled down by a factor of 0.9, and to create low and high pitched syllables in that voice, the fundamental frequency (F0) was scaled to either 77.8 or 155.6 Hz, respectively (half an octave above and below A2, 110Hz). In contrast, to create the “alto” voice of a primary school age boy the formants were scaled up by a factor of 1.4, and the F0s were set to 311 and 622 Hz (+/-0.5 octaves around A4) respectively to generate low and high pitched syllables in that voice.

To vary the perceived spatial location of the syllables, these morphed syllables were presented in a “virtual acoustic space” at a distance of 1 m, 60 degrees to the left or right off the midline in front of the person’s head, at eye level. To add realism, a small amount of reverberation was added to the sound by adding “specular reflections”, that is, each wall floor and ceiling of the room were treated as “sound mirrors” which will reflect sound essentially without frequency filtering but a flat reduction in amplitude. We chose an absorption coefficient of 0.6 for the virtual walls of this simple room model, an appropriate value to approximate the quite highly absorbent walls of the actual recording room. For a more detailed description of the present stimuli generation, see Retsa et al., (2018). E-prime software controlled stimulus delivery and recorded the subjects’ behavioural performance (www.pstnet.com/eprime).

### Procedure

Subjects performed 4 discrimination tasks, each comprising two separate blocks of trials varying on the to-be-discriminated dimension (Table 1); the order of blocks was counter-balanced across participants. Sounds were identical across blocks; only the specific instructions differed between blocks. Importantly, the sounds in all blocks were presented equi-probably to the right and the left side. Therefore, on each block, 50% of the stimuli were presented on the left side and 50% on the right side. Each block contained 160 trials and lasted approximately 5 minutes (i.e. 10 repetitions of each acoustic stimulus within a block). The subject’s task was to indicate as quickly and as accurately as possible whether the presented sound was i) presented on the left or right side - in the “spatial” task, ii) low or high pitch - in the “pitch” task, iii) a child or a man - in the “speaker” task, or iv) a ‘ti’ or a ‘ta’ - in the “syllable” task. The same response buttons (placed below the subject’s right index and right major finger) were used in all blocks, and their attribution was counterbalanced across participants.

**Table 1:** Task-relevant block type and features of stimuli that had to be discriminated

| Block type | Features to-be-discriminated |  |
| --- | --- | --- |
| Location | left side (50%) | right side (50%) |
| Pitch | low (50%) | high (50%) |
| Speaker | school age boy (50%) | adult man (50%) |
| Syllable | ti (50%) | ta (50%) |

### EEG recording and pre-processing

Continuous EEG was acquired at 1024Hz through a 128-channel Biosemi ActiveTwo AD-box (www.biosemi.com), referenced to the common mode sense (CMS; active electrode) and grounded to the driven right leg (DRL; passive electrode), which functions as a feedback loop driving the average potential across the electrode montage to the amplifier zero. Prior to epoching, the EEG was filtered (low-pass 40Hz; no high-pass; removed DC; 50Hz notch; using a second-order Butterworth filter with −12dB/octave roll-off that was computed linearly in both forward and backward directions to eliminate phase shifts). The EEG was segmented into peri-stimulus epochs spanning 100ms pre-stimulus to 500ms post-stimulus. Epochs with artefacts were rejected based on an automated artefact rejection criterion of ±80μV as well as visual inspection for eye blinks, eye movements and other sources of transient noise. Prior to group averaging, the data from electrodes with the highest proportion of artefacts were interpolated using 3-D splines (Perrin et al., 1987), for each subject separately. In addition, the data were baseline-corrected using the 100ms pre-stimulus period and recalculated against the average reference.

For each subject, eight sets of single-trial auditory evoked potentials (AEPs) were generated: one for the left-lateralised stimuli and one for the right-lateralised for each of the four tasks (i.e. location-left, location-right, pitch-left, pitch-right, speaker-left, speaker-right, syllable-left, syllable-right). Subsequently, these single-trial data from the four AEPs to left-lateralised stimuli were re-labelled so that electrodes over the left hemiscalp were treated as if they were located over the right hemiscalp and vice versa. In this way, data were always coded in terms of their contralaterality (contralateral versus ipsilateral to the presented sound location). The single-trial data from the left-lateralised and the right-lateralised stimuli were then averaged to generate the AEPs for each of the four tasks; the purpose was to assess the degree to which contralateral occipital activity is elicited by acoustically identical sounds as a function of task-relevant feature (location, pitch, speaker or syllable). Therefore, we refer to location, pitch, speaker and syllable conditions as well as contralateral and ipsilateral scalp sites with respect to stimuli. The average number (± SEM) of accepted EEG epochs for each subject and each of the above four conditions was 281±7, 285±6, 286±6 and 288±6, respectively. These values did not significantly differ (F(3,45) <1; p >0.75).

### ERP analyses

The ERP analyses followed closely the procedures employed in previous studies of the ACOP (Matusz et al., 2016; Feng et al., 2014; McDonald et al., 2013). In order to verify if the ACOP is elicited by all or just some types of task-relevant sounds, we first analysed the differences between contralateral and ipsilateral processing across the four conditions using mean voltages from ERPs over five selected parieto-occipital electrodes from each hemiscalp (see inset in Figure 2). These ERPs were analysed as a function of time using a 4 × 2 repeated-measures ANOVA with within-subject factors of Condition (four levels: location, pitch, speaker or syllable) and Contralaterality (two levels: contralateral, ispilateral). For this analysis we used an average reference as well as a temporal criterion for the detection of statistically significant effects (>10ms continuously at 1024Hz sampling rate) in order to correct for temporal auto-correlation at individual electrodes (Guthrie and Buchwald, 1991). This analysis allowed us to determine the onset latency of the ACOP in the current study.

**Figure 1.**
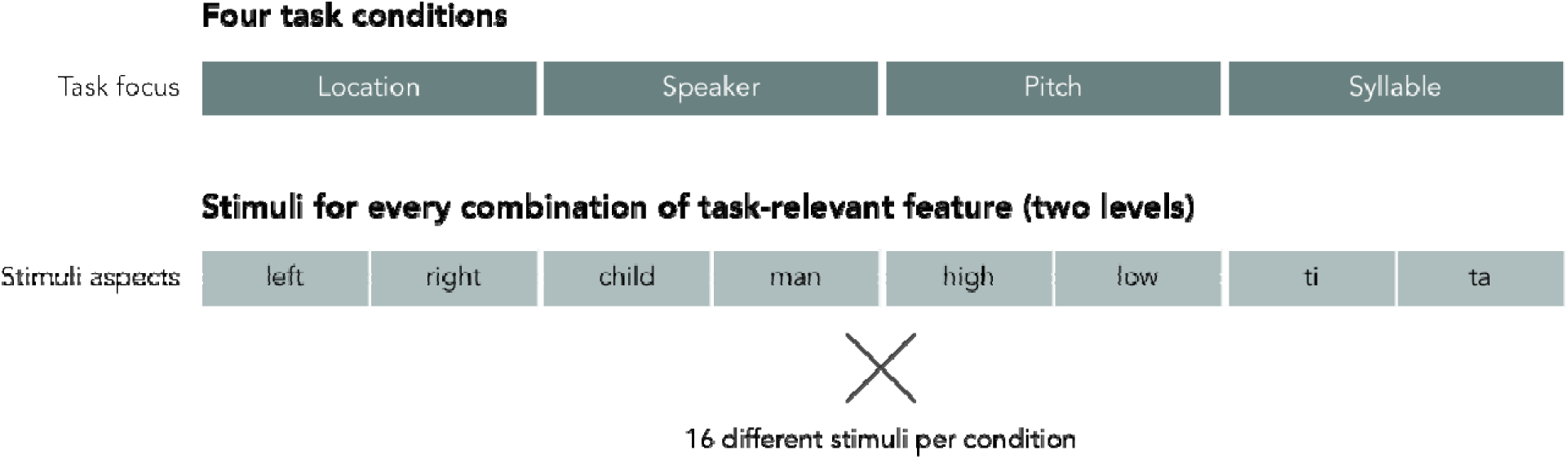
Schematic illustration of the task conditions and task-relevant features of the sound stimuli that had to be discriminated. Sound acoustics were held constant across blocks.

**Figure 2.**
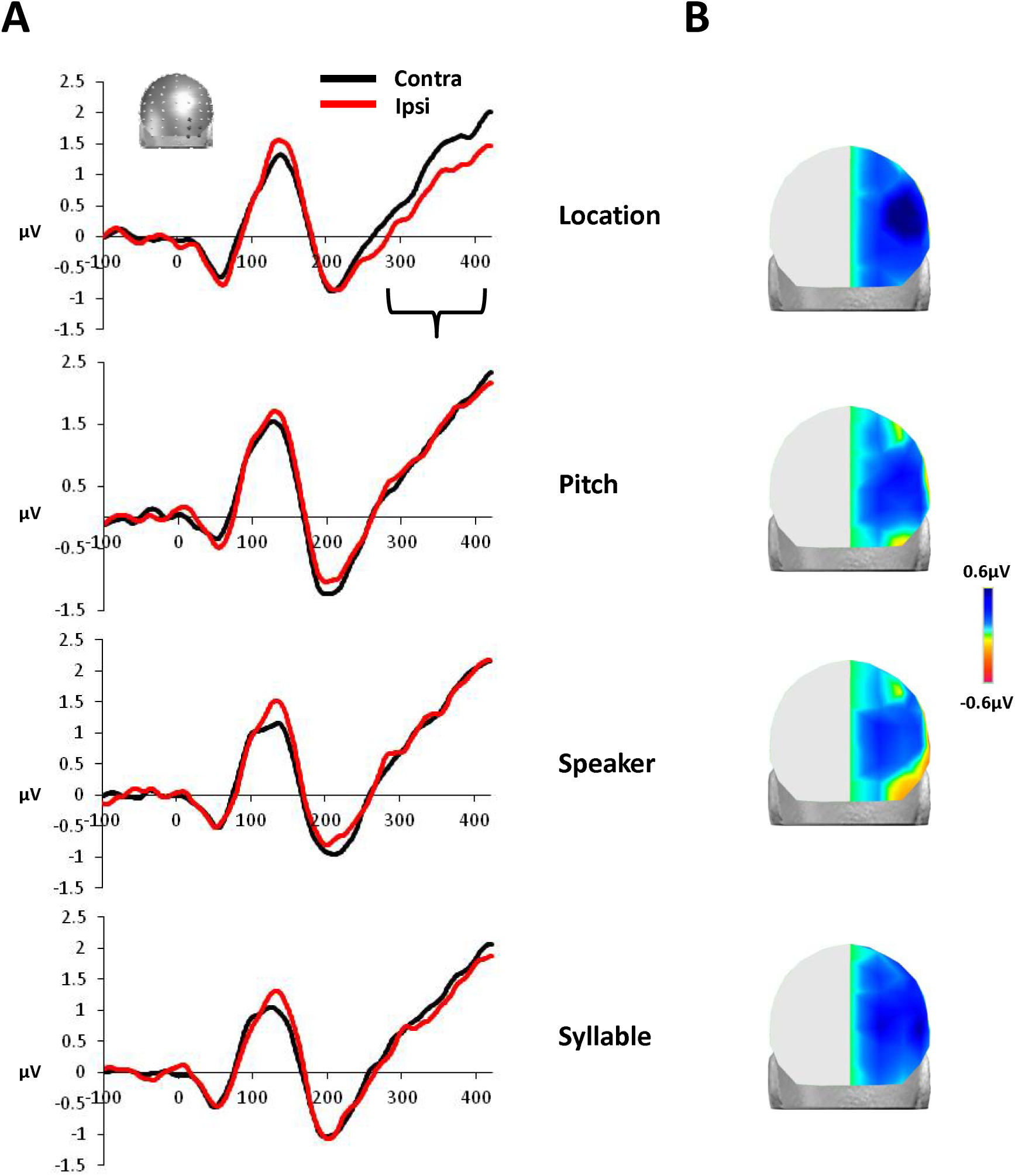
**A.** Contralateral and ipsilateral group-averaged ERPs, collapsed across 5 selected occipital electrodes (the inset depicts electrodes from the contralateral region of interest), plotted separately for each one of the four conditions. A reliable ACOP was observed only in the Location condition according to the results of the 2×4 repeated-measures ANOVA and the post-hoc t-test comparisons, over the 300-400ms post-stimulus period. **B.** The voltage topography quantified as the contralateral minus the ipsilateral difference amplitude over the 300-400ms time window is shown for each of the 4 conditions (projected onto the right hemisphere). The occipital positive voltages in response to the sounds appear to be clearly enhanced on the contralateral side, but only in the Location condition.

Next, we aimed to provide direct evidence that contexts where spatial location is task-relevant lead to a reliable and larger ACOP, as compared to when other sound dimensions are task-relevant. Finally, we sought to determine whether any observed ACOP was the result of amplitude enhancement over the contralateral scalp and/or suppression over the ipsilateral scalp. To this end, we analysed mean ERP amplitude over the 300-400ms post-sound time-window in a 4×2 rmANOVA with within-subject factors of Condition (four levels: location, pitch, speaker or syllable) and Contralaterality (two levels: contralateral, ispilateral). As detailed below, in the face of a reliable 2-way interaction we conducted follow-up ANOVAs to address the specific questions regarding task-selective presence of the ACOP and the nature (enhancement/suppression) of the underlying brain activity.

### Source estimations

We estimated the likely underlying intracranial sources of the ERP effects identified in the above-mentioned scalp measurements using a distributed linear inverse solution (minimum norm) combined with the LAURA (local autoregressive average) regularisation approach (Grave de Peralta Menendez et al., 2001, 2004; see also Michel et al., 2004 for a review). LAURA selects the source configuration that best mimics the biophysical behaviour of electric vector fields (i.e., activity at one point depends on the activity at neighbouring points according to electromagnetic laws). In our study, as part of the regularisation strategy, homogenous regression coefficients in all directions and within the whole solution space were used. The solution space was calculated on a realistic head model that included 3005 nodes, selected from a grid equally distributed within the grey matter of the Montreal Neurological Institute’s average brain (courtesy of Grave de Peralta Menendez and Gonzalez Andino). The head model and lead field matrix were generated within the Spherical Model with Anatomical Constraints (SMAC; Spinelli et al., 2000 as implemented in Cartool (Brunet et al., 2011)). As an output, LAURA provides current density measures; their scalar values were evaluated at each node. The statistical significance criterion at an individual solution point was set at p<0.05. Only clusters with at least 15 contiguous significant nodes were considered reliable in an effort to correct for multiple comparisons, and this size was based on randomisation thresholds determined with Alphasim software (see also Toepel et al., 2009; De Lucia et al., 2012; Knebel and Murray, 2012; Thelen et al. 2014; Matusz et al., 2015b; Matusz et al., 2016 for similar implementations). The source estimations of the mean ACOP were calculated for each subject and condition over the time-period exhibiting a reliable Condition × Laterality interaction in the ERP analysis described above. Statistical analysis of source estimations was performed on the difference between contralateral and ipsilateral AEPs (one-way ANOVA with the within-subjects 4-level factor of Condition).

### Results

Behavioural results on this task appear in Retsa et al. (2018). Briefly, subjects performed at near-ceiling levels (i.e. >90%). AEP voltage waveforms averaged across a set of 5 parieto-occipital electrodes (inset of Figure 2) were submitted to a 2 x 4 repeated-measures ANOVA as a function of time. There were reliable main effects of Condition (90–191ms and 280–400ms) and Contralaterality (118–161ms and 328–400ms). In addition, there was a reliable two-way interaction between Condition and Contralaterality over the 300–400ms post-stimulus time window. Post-hoc pairwise t-tests showed significant differences between contralateral and ipsilateral responses (and therefore, the presence of the ACOP), only for the Location condition. In the three other conditions (Pitch, Speaker, Syllable), no significant differences between contralateral and ipsilateral responses were observed. Figure 2A displays contralateral and ipsilateral ERPs averaged over the five electrodes for each of the four different task conditions. Figure 2B displays the voltage topography for the four different conditions, quantified as the contralateral minus ipsilateral difference amplitude over the 300–400ms time window (projected onto the right hemisphere).

Figure 3 shows the mean amplitudes of the ACOP (calculated as the difference between contralateral and ipsilateral responses over the 300–400ms period) for each condition (Figure 3A), as well as the mean amplitude of contralateral and ipsilateral responses plotted separately for each condition (over the same 300–400ms time window) (Figure 3B). The two-way ANOVA revealed a main effect of Contralaterality (F_(1,15)_=4.83; p=0.044; η_p_^2^=0.24) that was further modulated by Condition (F_(1.86,27.94)_=11.09; p<0.001; η_p_^2^=0.43). Given this interaction, we next tested if the ACOP was reliable for each condition. This was achieved with a series of paired t-tests comparing contralateral versus ipsilateral responses for each condition separately. This comparison was significant only for the Location condition (t_(15)_=3.6; p<0.01). All other conditions were not reliable (p>0.24 in all cases). Next, we assessed if the ACOP was larger in the Location than all other conditions, comparing the mean ACOP (contralateral minus ipsilateral response difference). The ACOP magnitude was larger for the Location condition as compared to Pitch (t_(15)_=3.99; p<0.001), Speaker (t_(15)_=3.8; p<0.002), as well as Syllable (t_(15)_=3.3; p<0.005). No significant differences between the other dimensions were observed (all p’s >0.11). Finally, we tested the nature (enhancement/suppression) of the underlying brain activity via two 1-way ANOVAs: one on the ipsilateral responses over the 300-400ms period and one on the respective contralateral responses with Condition as the within-subject factor. The ANOVA on the ipsilateral responses showed a main effect of Condition (F_(3,45)_=4.63; p=0.007; η_p_^2^=0.24), whereas that on the contralateral responses did not (F_(2.06,30.91)_<1; p>0.77; η_p_^2^=0.02). Post-hoc t-tests (Holm-Bonferroni correction applied) showed significant differences in the ipsilateral responses between Location and Pitch (t_(15)_=2.92; p=0.01) and Location and Speaker (t_(15)_=3.78; p=0.002). None of the other results were statistically reliable (p’s>.12).

**Figure 3.**
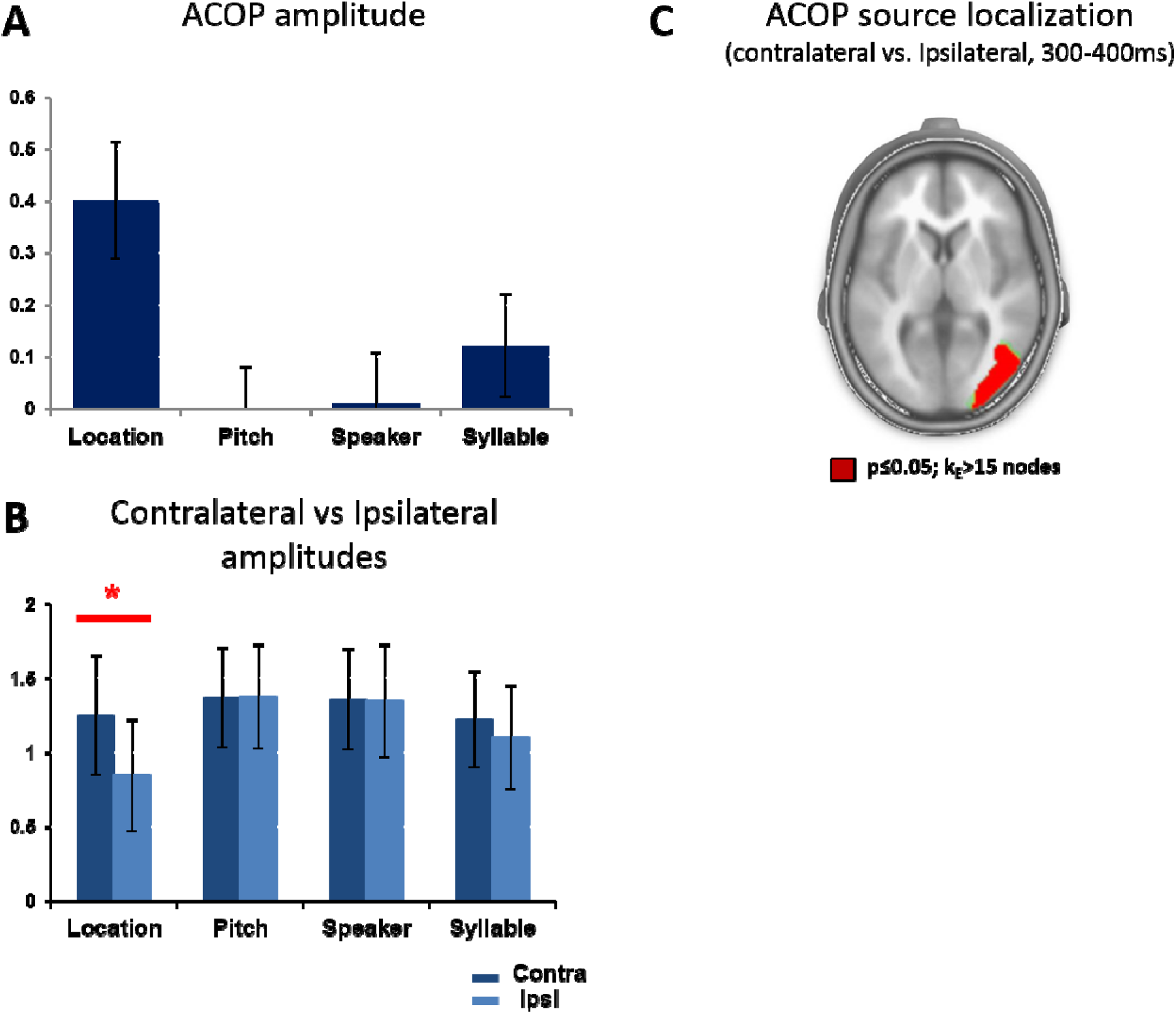
A. Mean ACOP difference amplitudes (contralateral minus ipsilateral) over the 300-400 ms time period averaged over the 5 occipital electrodes for each of the 4 different conditions. ACOP was significantly larger for the Location condition compared to either Pitch, Speaker or Syllable. B. Mean amplitude of contralateral and ipsilateral responses plotted separately for each condition over the 300-400 ms time-period. T-tests between contralateral and ipsilateral responses was only significant for the Location condition (p<.01). ACOP in the Location condition appears as a result of a suppression of the ipsilateral responses rather than an enhancement of the contralateral ones. C. Differential source activity across conditions was observed in the lateral occipital cortex (a representative axial slice is displayed) with stronger activity found for the Location condition compared to any other condition. Symmetrical activity was computed across the two hemispheres for the purposes of the source localisation (in a manner akin to what was done with the surface ERP data). Consequently, we plot sources only in the right hemisphere.

Source estimations were then calculated for the difference (contralateral minus ipsilateral) AEPs from the four conditions over the 300–400ms post-stimulus period (i.e. the time-period that exhibited a significant ACOP as measured at the scalp surface). The one-way four-level ANOVA revealed a significant main effect of Condition on the estimated source activity. Differential source activity was localised to clusters within the lateral occipital cortex (local maximum F-value at 35, −75, 11mm using the Talairach and Tournoux (1988) atlas) (Figure 3C). This locus is in line with previous research (cf., Figure 2c in Matusz et al., 2016; also McDonald et al., 2013; Feng et al., 2014). Post-hoc t-tests on the differential activity at the local maximum showed stronger activity in the Location condition compared to Pitch (t_(15)_=3.5; p<0.01) and Syllable (t_(15)_=3.8; p<0.001), and a trend for stronger activity compared to Speaker (t_(15)_=1.9; p=0.064).

### Discussion

Laterally presented sounds can elicit activations of the contralateral occipital cortex around 300ms post-stimulus onset. To investigate the effect of selective attention to specific sound features on the presence of contralateral cross-modal activations, we compared the occipital brain responses during the discrimination of sounds across four different dimensions: location, pitch, syllable type, and speaker identity. Participants were presented with an identical set of sounds on every block of trials; the only parameter varying was the dimension that was relevant to the discrimination task at-hand. Our results demonstrated that the ACOP can be elicited by task-relevant, real-world like sounds. However, the same sounds elicited the ACOP only when participants had to discriminate the location of sounds. In all the other types of blocks of trials, no reliable cross-modal activation of visual cortices was observed. Therefore, we provide evidence that when the auditory stimuli are task-relevant, the presence of differential contralateral vs. ipsilateral sound-elicited activation of visual cortices depends on the spatial dimension of sound processing being task-relevant.

The timing of the activation of visual cortices induced by lateralised sounds in the present study (over the 300–400ms post-stimulus time window) is consistent with the latency of the ACOP observed in previous studies (Hillyard et al., 2016; Matusz et al., 2016). Furthermore, the sources of this activity within the lateral occipital cortex were found in loci similar to those in these previous studies. Most importantly, the results of Matusz et al. (2016) indicated that the ACOP is not an automatic process, as context – specifically, stimulus regularities – determines its presence. When sounds are task-irrelevant, such as in the case of a passive auditory task, the ACOP depends on the unpredictability of the sound location in space. Specifically, the ACOP was observed only when the location of the task-irrelevant sounds was unpredictable. When the location of those stimuli was implicitly predictable, then the contralateral occipital response was suppressed and no ACOP was observed. The absence of the ACOP in that case was hypothesised to be the result of an effective inhibition of the processing of completely task-irrelevant sounds. In contrast, when sounds were presented equiprobably on the left and on the right sides, the inhibition of exogenous shift of attention (which is considered to be the basis of the ACOP) was not possible. The present results extend these findings to indicate that when voluntary control of selective attention is at play, spatial unpredictability alone is insufficient for the ACOP to manifest.

In our study, all conditions involved sounds that were spatially unpredictable (at least between left and right sides). However, the ACOP was elicited only when location was the task-relevant dimension. The present finding of task-selective cross-modal activations of visual cortices by sounds is consistent with a recent ERP study that compared spatial and temporal processing of lateralised sounds (Campus et al., 2017). They observed responses within the visual cortex, including during the ACOP time window, only when the location, but not timing, of the auditory stimuli was task-relevant. Cross-modal activations of occipital cortices by sounds were observed only when a spatial, but not temporal, bisection task was employed, and despite the fact that identical stimuli were used in both cases. Their data provided further evidence for the involvement of visual cortex in the perception of space and its potential role in the construction of a spatial metric. Other, fMRI data would however suggest that auditory activations of visual cortices critically depend on participants selectively attending to the auditory (versus visual) modality (Cate et al., 2009; see also Laurienti et al., 2002). However, a straightforward comparison with the present results, and studies of the ACOP more generally, is hampered by the fact that the paradigm of Cate et al. (2009) entailed blocked lateralisation of sounds as well as presentation of multisensory stimuli to which participants were instructed to attend with respect to the visual or auditory feature. Thus, there was no variation in the location of the stimuli within a block of trials, but rather only across blocks. By contrast, studies of the ACOP suggest that cross-modal activation of visual cortices is related to spatial processing in one form or another (e.g. either attending to spatial locations and/or explicitly processing the spatial information within the stimuli). Previous studies have shown that vision plays an important role in auditory spatial processing (Driver & Spence, 2004), and one proposition is that spatial attention processes may thus be dominated by visual representations of location, based on the contention of superior spatial acuity in vision than audition (Welch and Warren, 1980). It remains unresolved, however, to what extent spatial representations are truly “visual”, even if localised within nominally visual cortices, or instead are multisensory both in their content and functional consequences on perception and behaviour (see Ten Oever et al., 2016 for discussion as well as Eimer et al., 2002; Gamble & Luck, 2011). A further possibility is that cross-modal lateralised ERP differences at the occipital scalp and in visual cortices are instead a result of a carryover effect of lateralised processing of contralateral vs. ipsilateral sound stimuli occurring first in auditory cortices, which is subsequently modulated by top-down attentional control mechanisms (Plass et al., 2019). Such an account is (at least partially) supported by evidence of spatial attention effects on early-latency lateralised ERP components presumed to originate within auditory cortices (Hillyard et al., 1973; Alho et al., 1999). However, studies of the ACOP to date would suggest that its elicitation is not a simple carry-over effect of responses within auditory cortices, but rather a more direct impact of auditory information on multisensory spatial representations within visual cortices (reviewed in Hillyard et al., 2016).

Interestingly, in the current study, the ACOP seems to be the result of reduced activity within the ipsilateral hemisphere (see Figure 3B) instead of enhanced contralateral activity, as reported in prior investigations of the ACOP (McDonald et al., 2013; Feng et al., 2014; Matusz et al., 2016). The task-relevance of the stimuli appears to be a crucial difference across studies. In the previous cases, where an enhanced occipital response contralaterally to the sound location was elicited, the lateralised sounds were either uninformative cues or part of a passive task, and the ACOP was interpreted as reflecting shifts of exogenous, involuntary spatial attention. Here, the lateralised sounds were part of the task and were actively anticipated, which involves top-down, goal-based orienting of spatial attention towards the sounds. When voluntary spatial attention is directed towards the location aspect of the sounds then potentially suppressive processes are implicated; specifically, our results suggests that the irrelevant side of space is suppressed and, as a consequence, the ACOP is observed. In contrast, when attention is focused on non-spatial aspects of the sounds (such as pitch, speaker identity, and syllable type) no ACOP is observed. This account is corroborated by the findings of Matusz et al. (2016) in the condition where the location of the irrelevant sounds was predictable and no ACOP was observed. Matusz et al. (2016) interpreted this result in terms of top-down suppression of contralateral cross-modal enhancements. In the present results, focusing on non-spatial aspects when sounds are actively anticipated may also result in a suppression of these contralateral cross-modal responses.

The present findings shed light on the necessary conditions for cross-modal activation of visual cortices by lateralised sounds (ACOP) (see Figure 4 for a summary). The ACOP depends on both stimulus regularities and task-relevance; evidence that further supports its non-automatic nature. When the lateralised sounds are task-relevant, then ACOP is elicited only when the location aspect of the sounds is currently important, highlighting the role visual cortices (and likely visual processes) play in auditory spatial functions. In contrast, when the top-down attention of participants is focused on non-location aspects, cross-modal activation of visual cortex is not observed. These results suggest preferential cross-modal interactions for spatial processing. In conclusion, we demonstrate the interplay between task-relevance and spatial unpredictability in producing the cross-modal activation of visual cortices.

**Figure 4.**
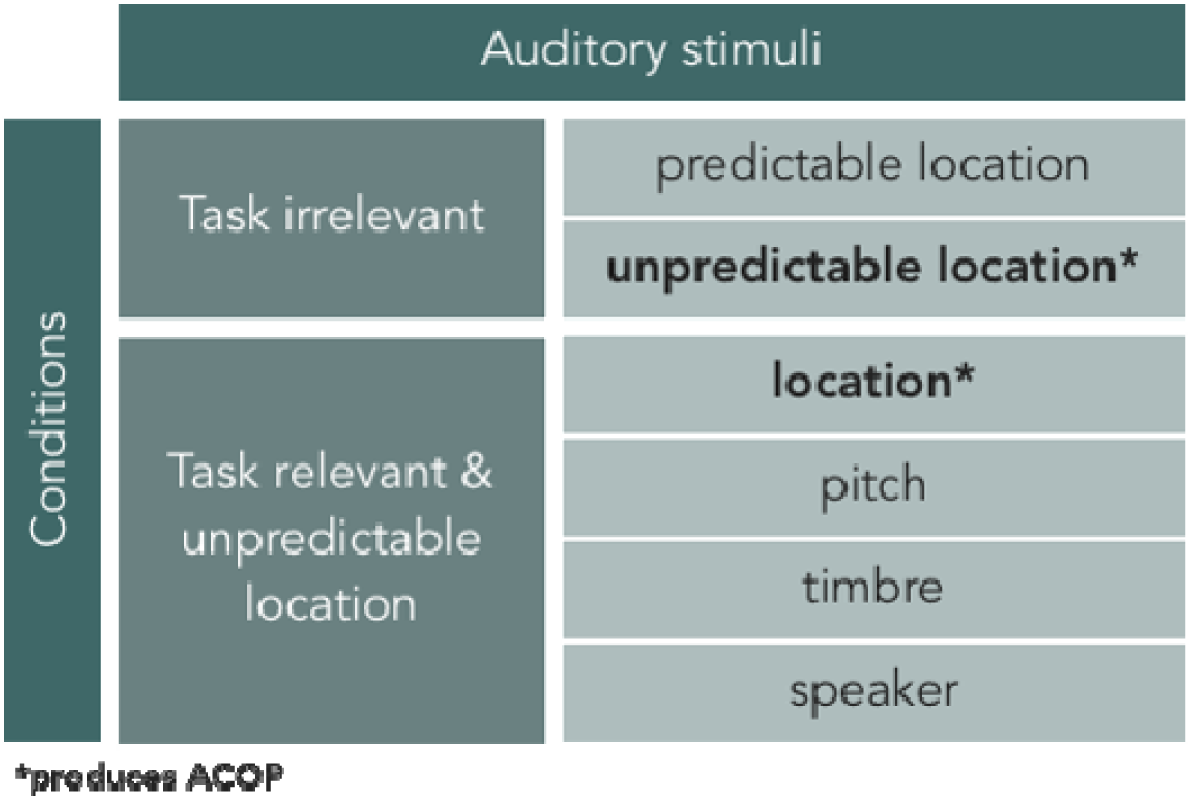
Schematic illustration of the conditions under which ACOP is produced and the conditions under which ACOP is not observed.

## Acknowledgement

Financial support was provided by the Swiss National Science Foundation (grants: 320030_149982 and 320030_169206 to M.M.M., PZ00P1_174150 to P.J.M., and the National Centre of Competence in research project “SYNAPSY, The Synaptic Bases of Mental Disease” [project 51AU40_125759]). P.J.M. receives support from the Pierre Mercier Foundation. P.J.M. and M.M.M. are both supported by Fondation Asile des aveugles. Also, this work has been supported by a “Royal Society International Exchange Project support grant (to J.S. and M.M.M.). These sources had no further role in this study design, in the data collection and analysis, in the writing of the report, and in the decision to submit the paper for publication.

## References

Alho, K., Medvedev, S.V., Pakhomov, S.V., Roudas, M.S., Tervaniemi, M., Reinikainen, K., Zeffiro, T., Naatanen, R., 1999. Selective tuning of the left and right auditory cortices during spatially directed attention. Brain Res Cogn Brain Res., 7, 335–341.

Beer, A.L., Plank, T. & Greenlee, M.W., 2011. Diffusion tensor imaging shows white matter tracts between human auditory and visual cortex. Exp. Brain Res., 213, 299–308.

Bolognini, N., Senna, I., Maravita, A., Pascual-Leone, A., & Merabet, L.B., 2010. Auditory enhancement of visual phosphene perception: the effect of temporal and spatial factors and of stimulus intensity. Neurosci. Lett., 477(3), 109–114.

Brang, D., Towle, V.L, Suzuki, S., Hillyard, S.A., Di Tusa, S., Dai, Z., Tao, J., Wu, S., & Grabowecky, M., 2015. Peripheral sounds rapidly activate visual cortex: evidence from electrocorticography. J. Neurophysiology, 114(5), 3023–3028.

Brunet, D., Murray, M.M., Michel, C.M., 2011. Spatiotemporal analysis of multichannel EEG: CARTOOL. Comput. Intell. Neurosci. doi:10.1155/2011/813870

Budinger, E., Heil, P., Hess, A., Scheich, H., 2006. Multisensory processing via early cortical stages: connections of the primary auditory cortical field with other sensory systems. Neuroscience, 143, 1065–1083.

Campus, C., Sandini, G., Morrone, C. M., Gori, M., 2017. Spatial localization of sound elicits early responses from occipital visual cortex in humans. Scientific Reports, 7, 10415, doi:10.1038/s41598-017-09142-z

Cappe, C., Thut, G., Romei, V., Murray, M.M., 2010. Auditory-visual multisensory interactions in humans: timing, topography, directionality, and sources. J. Neurosci., 30(38), 12572–12580.

Cappe, C., Barone, P., 2005. Heteromodal connections supporting multisensory integration at low levels of cortical processing in the monkey. Eur. J. Neurosci., 22, 2886–2902.

Cate, A.D., Herron, T.J., Yund, E.W., Stecker, G.C., Rinne, T., Kang, X., Petkov, C.I., Disbrow, E.A., Woods, D.L. 2009. Auditory attention activates peripheral visual cortex. PLoS One, https://doi.org/10.1371/journal.pone.0004645

Clavagnier, S., Falchier, A., Kennedy, H., 2004. Long-distance feedback projections to area V1: implications for multisensory integration, spatial awareness, and visual consciousness. Cogn. Affect. Behav. Neurosci., 4, 117–126.

De Lucia, M., Tzovara, A., Bernasconi, F., Spierer, L., Murray, M.M., 2012. Auditory perceptual decision-making based on semantic categorization of environmental sounds. Neuroimage, 60, 1704–1715.

De Meo, R., Murray, M.M., Clarke, S., Matusz, P.J., 2015. Top-down control and early multisensory processes: chicken vs. egg. Front. Integr. Neurosci. 9, 17. http://dx.doi.org/10.3389/fnint.2015.00017.

Driver, J., Spence, C., 2004. Cross-modal spatial attention: evidence from human performance. In: Spence C, Driver J, editors. Cross-modal space and cross-modal attention. Oxford University Press, Oxford, UK, 179–220.

Eckert, M. A., Kamdar, N. V., Chang, C. E., Beckmann, C. F., Greicius, M. D., & Menon, V., 2008. A cross-modal system linking primary auditory and visual cortices: evidence from intrinsic fMRI connectivity analysis. Human brain mapping, 29(7), 848–857.

Eimer, M., van Velzen, J., Driver, J., 2002. Cross-modal interactions between audition, touch and vision in endogenous spatial attention: ERP evidence on preparatory states and sensory modulations. J. Cogn. Neurosci., 14(2), 254–271.

Falchier, A., Schroeder, C. E., Hackett, T. A., Lakatos, P., Nascimento-Silva, S., Ulbert, I.,…Smiley, J. F., 2010. Projection from visual areas V2 and prostriata to caudal auditory cortex in the monkey. Cerebral cortex, 20(7), 1529–1538.

Falchier, A., Clavagnier, S., Barone, P., Kennedy, H., 2002. Anatomical evidence of multimodal integration in primate striate cortex. J Neurosci., 22, 5749–5759.

Feng, W., Stllrmer, V.S., Martinez, A., McDonald, J.J., Hillyard, S.A., 2014. Sounds activate visual cortex and improve visual discrimination. J. Neurosci, 34(29), 9817–9824.

Gamble, M. L., & Luck, S. J., 2011. N2ac: an ERP component associated with the focusing of attention within an auditory scene. Psychophysiology, 48(8), 1057–1068. doi:10.1111/j.1469-8986.2010.01172.x Giard, M.H., Peronnet, F., 1999. Auditory-visual integration during multimodal object recognition in humans: a behavioral and electrophysiological study. J Cogn. Neurosci., 11, 473–490.

Gleiss, S, Kayser, C., 2014. Acoustic noise improves visual perception and modulates occipital oscillatory states. J. Cogn. Neurosci., 26, 699–711.

Grave de Peralta Menendez, R., Gonzalez Andino, S., Lantz, G., Michel, C.M., Landis, T., 2001. Noninvasive localization of electromagnetic epileptic activity. I. Method descriptions and simulations. Brain Topogr. 14, 131–137.

Grave de Peralta Menendez, R., Murray, M.M., Michel, C.M., Martuzzi, R., Gonzalez Andino, S.L., 2004. Electrical neuroimaging based on biophysical constraints. NeuroImage 21, 527–539. doi:10.1016/j.neuroimage.2003.09.051

Guthrie, D., Buchwald, J.S., 1991. Significance testing of difference potentials. Psychophysiology 28, 240–244.

Hillyard, S.A., Stllrmer, V.S., Feng, W., Martinez, A., McDonald, J.J. 2016. Cross-modal orienting of visual attention. Neuropsychologia, 83, 170–178,

Hillyard, S.A., Hink, R.F., Schwent, V.L., Picton, TW., 1973. Electrical signs of selective attention in the human brain. Science, 182(4108), 177–180.

Ives, D. T., Smith, D. R., Patterson, R. D. 2005. Discrimination of speaker size from syllable phrases, J. Acoust. Soc. Am. 118, 3816.

Kayser, S.J., Kayser, C., 2018. Trial by trial dependencies in multisensory perception and their correlates in dynamic brain activity. Scientific reports, 8(1), 3742.

Kayser, S.J., Philiastides, M.G., Kayser, C., 2017. Sounds facilitate visual motion discrimination via the enhancement of late occipital visual representations. Neuroimage, 148, 31–41.

Knebel, J.F., Murray, M.M., 2012. Towards a resolution of conflicting models of illusory contour processing in humans. Neuroimage, 59, 2808–2817,

Laurienti, P.J., Burdette, J.H., Wallace, M.T., Yen, Y.F., Field, A.S., Stein, B.E., 2002. Deactivation of sensory-specific cortex by cross-modal stimuli. J. Cogn. Neurosci. 14, 420–429.

Leo, F., Romei, V., Freeman, E., Ladavas, E., Driver, J., 2011. Looming sounds enhance orientation sensitivity for visual stimuli on the same side as such sounds. Exp.Brain Research, 213(2-3), 193–201.

Lovelace, C. T., Stein, B. E., & Wallace, M. T., 2003. An irrelevant light enhances auditory detection in humans: A psychophysical analysis of multisensory integration in stimulus detection. Cognitive Brain Research, 17, 447–453.

McDonald, J.J., Störmer, V.S, Martinez, A., Feng, W., Hillyard, S.A., 2013. Salient sounds activate human visual cortex automatically. J. Neurosci., 33(21), 9194–9201.

Matusz, P. J., Turoman, N., Tivadar, R. I., Retsa, C., & Murray, M. M., 2019a. Brain and cognitive mechanisms of top–down attentional control in a multisensory world: Benefits of electrical neuroimaging. Journal of cognitive neuroscience, 31(3), 412–430.

Matusz, P. J., Merkley, R., Faure, M., & Scerif, G., 2019b. Expert attention: Attentional allocation depends on the differential development of multisensory number representations. Cognition, 186, 171–177.

Matusz, P. J., Dikker, S., Huth, A. G., & Perrodin, C., 2019c. Are we ready for real-world neuroscience?.Journal of Cognitive Neuroscience, 31, 327–338.

Matusz, P.J., Retsa, C., Murray, M.M., 2016. The context-contingent nature of cross-modal activations of the visual cortex. Neuroimage, 125, 996–1004.

Matusz, P. J., Broadbent, H., Ferrari, J., Forrest, B., Merkley, R., & Scerif, G., 2015a. Multi-modal distraction: Insights from children’s limited attention. Cognition, 136, 156–165.

Matusz, P.J., Thelen, A., Amrein, S., Geiser, E., Anken, J., Murray, M.M., 2015b. The role of auditory cortices in the retrieval of single-trial auditory-visual object memories. European J. Neurosci., 41, 699–708.

Matusz, P. J., Thelen, A., Amrein, S., Geiser, E., Anken, J., & Murray, M. M., 2015c. The role of auditory cortices in the retrieval of single-trial auditory–visual object memories. European Journal of Neuroscience, 41(5), 699–708.

Matusz, P.J., Eimer, M., 2013. Top-down control of audiovisual search by bimodal search templates. Psychophysiology 50, 996–1009.

Matusz, P.J., Eimer, M., 2011. Multisensory enhancement of attentional capture in visual search. Psychol. Bull. Rev. 18, 904–909.

McDonald, J.J., Teder-Sälejärvi, W.A., Hillyard, S.A., 2000. Involuntary orienting to sound improves visual perception. Nature 407, 906–908.

Mercier, M.R., Foxe, J.J., Fiebelkorn, I.C., Butler, J.S., Schwartz, T.H., & Molholm, S. 2013. Auditory-driven phase reset in visual cortex: Human electrocorticography reveals mechanisms of early multisensory integration. Neuroimage, 79, 19–29.

Michel, C.M., Murray, M.M., Lantz, G., Gonzalez, S., Spinelli, L., Grave de Peralta, R., 2004. EEG source imaging. Clin. Neurophysiol. 115, 2195–2222.

Molholm, S., Ritter, W., Murray, M.M., Javitt, D.C., Schroeder, C.E., Foxe, J.J., 2002. Multisensory auditory-visual interactions during early sensory processing in humans: a high-density electrical mapping study. Cogn. Brain Research, 14(1), 115–128.

Murray, M.M., Thelen, A., Thut, G., Romei, V., Martuzzi, R., Matusz, P.J., 2016. The multisensory function of the human primary visual cortex. Neuropsychologia, 83, 161–169.

Odgaard, E.C., Arieh, Y., Marks, L.E., 2004. Brigher noise: Sensory enhancement of perceived loudness by concurrent visual stimulation. Cog., Aff., & Behav. Neurosci., 4(2), 127–132.

Oldfield, R.C., 1971. The assessment and analysis of handedness: the Edinburgh inventory. Neuropsychologia, 9(1), 97–113.

Perrin, F., Pernier, J., Bertrand, O., Giard, M.H., Echallier, J.F., 1987. Mapping of scalp potentials by surface spline interpolation. Electorencephalogr. Clin. Neurophysiol., 66, 75–81.

Plass, J., Ahn, E., Towle, V.L., Stacey, W.C., Wasade, V.S., Tao, J., Wu, S., Issa, N.P., Brang, D., 2019. Joint encoding of auditory timing and location in visual cortex. J. Cogn. Neurosci., 31(7), 1002–1017.

Raij, T., Uutela, K., Hari, R., 2000. Audiovisual integration of letters in the human brain. Neuron, 28, 617–625.

Raij, T., Ahveninen, J., Lin, F.H., Witzel, T., Jääskeläinen, I.P., Letham, B., … Belliveau, J.W., 2010. Onset timing of cross-sensory activations and multisensory interactions in auditory and visual sensory cortices. Eur. J. Neurosci. 31, 1772–1782.

Rockland, K.S., Ojima, H., 2003. Multisensory convergence in calcarine visual areas in macaque monkey. Int. J. Psychophysiol., 50, 19–26.

Romei, V., Murray, M.M., Cappe, C., Thut, G., 2013. The contributions of sensory dominance and attentional bias to cross-modal enhancement of visual cortex excitability. J. Cogn. Neurosci., 25, 1122–1135.

Romei, V., Murray, M.M., Cappe, C., Thut, G., 2009. Preperceptual and stimulus-selective enhancement of low-level human visual cortex excitability by sounds. Curr. Biol. 19, 1799–1805.

Romei, V., Murray, M.M., Merabet, L.B., Thut, G., 2007. Occipital transcranial magnetic stimulation has opposing effects on visual and auditory stimulus detection: implications for multisensory interactions. J. Neurosci. 27, 11465–11472.

Spierer, L., Manuel, A.L., Bueti, D., Murray, M.M., 2013. Contributions of pitch and bandwidth to sound-induced enhancement of visual cortex excitability in humans. Cortex 49, 2728–2734.

Spinelli, L., Andino, S.G., Lantz, G., Seeck, M., Michel, C.M., 2000. Electromagnetic inverse solutions in anatomically constrained spherical head models. Brain Topogr. 13, 115–125.

Störmer, V.S., McDonald, J.J., Hillyard, S.A., 2009. Cross-modal cueing of attention alters appearance and early cortical processing of visual stimuli. Proc. Natl. Acad. Sci. U. S. A. 106, 22456–22461.

Retsa, C.*, Matusz*, P.J, Schnupp, J.W.H., Murray, M.M., 2018. What’s what in auditory cortices? Neuroimage, 176, 29–40.

Ten Oever, S., Romei, V., van Atteveldt, N., Soto-Faraco, S., Murray, M.M., Matusz, P.J., 2016. The COGs (context, object, and goals) in multisensory processing. Exp. Brain Research, 234(5), 1307–1323.

Thelen, A.*, Matusz, P.J.*, Murray, M.M., 2014. Multisensory context portends object memory. Current Biology 24(16), R734–735.

Tivadar, R.I., Retsa, C., Turoman, N., Matusz, P.J., Murray, M.M., 2018. Sounds enhance visual completion processes. Neuroimage, 179, 480–488.

Toepel, U., Knebel, J.F, Hudry, J., le Coutre, J., Murray, M.M., 2009. The brain tracks the energetic value in food images. Neuroimage, 44, 967–974.

Wang, Y., Celebrini, S., Trotter, Y., Barone, P., 2008. Visuo-auditory interactions in the primary visual cortex of the behaving monkey: electrophysiological evidence. BMC Neurosci., 9:79.

Welch, R.B., Warren, D.H., 1980. Immediate perceptual response to intersensory discrepancy. Psychol. Bull., 88(3), 638–667.

